# Automated SP3 Workflow Enables Robust Sample Preparation on Proteonano Ultraplex Proteomics Platform

**DOI:** 10.1101/2025.05.27.656469

**Authors:** Yi Wang, Libing Wang, Yonghao Zhang, Xiehua Ouyang, Hao Wu

## Abstract

The efficiency and reproducibility of sample preparation are critical prerequisites for obtaining reliable results in bottom-up proteomics. To conclude, we systematically evaluated three commonly used sample preparation methods: automated single-pot solid-phase-enhanced sample preparation (SP3), precipitation by organic solvents, and filter-aided sample preparation method (FASP). When processing varying input amounts of HeLa cell lysate (10, 20, 50, 100, and 150 μg), the SP3 method demonstrated strong linearity (R^2^ = 0.95), indicating excellent quantitative consistency across a broad protein input range. A comparison between manual SP3 and manual precipitation protocols revealed comparable protein identification numbers, with a correlation coefficient of approximately 0.97. We further developed an automated version of the SP3 method based on the Proteonano Ultraplex Proteomics platform (autoSP3), which yielded highly consistent results with the manual approach, achieving a protein identification overlap of 98.5% (6,949 out of 7,056). These findings support the suitability of the autoSP3 workflow as a robust and scalable approach for high-throughput and reproducible sample preparation in proteomics sample preparation.

## Introduction

Proteins, as central executors of biological processes, orchestrate intricate physiological and pathological mechanisms through dynamic molecular interactions. Recent advancements in proteomics have revolutionized our capacity for system-level protein characterization, where efficient sample preparation remains the critical gateway to high-fidelity analyses^1^. Conventional protein extraction strategies, while foundational, face insurmountable challenges including procedural complexity, significant analyte loss, and detergent interference - limitations that increasingly constrain modern high-throughput proteomic investigations^1^.

The advent of single-pot solid-phase-enhanced sample preparation (SP3) technology marks a paradigm shift in protein enrichment methodologies. This innovative approach harnesses functionalized magnetic nanoparticles that reversibly bind proteins through solvent-mediated interfacial interactions. Subsequent aqueous-phase elution coupled with on-bead digestion achieves detergent-free protein recovery with exceptional yield and reproducibility^2,3^.

Mechanistically distinct from traditional methods, SP3 eliminates multiple liquid transfer steps through its single-tube workflow, effectively minimizing sample loss and cross-contamination risks. Its versatility across diverse biological matrices – from cellular lysates to complex biofluids – positions SP3 as a universal platform for next-generation proteomic studies.

Significantly, SP3 technology demonstrates transformative potential for automated proteomic workflows. The escalating demands of clinical proteomics necessitate robust, standardized sample processing platforms. Recent integration of SP3 with commercial automation platforms has achieved unprecedented throughput in low-input clinical specimens^3^. Pioneering work by Müller *et al.* ^3^ at the German Cancer Research Center exemplifies this synergy, combining automated SP3 with Covaris AFA-based acoustic shearing to establish a closed-system pipeline that reduces manual intervention by 70% while enhancing inter-laboratory reproducibility.

To address the above challenges, this study presents an automated SP3-based sample preparation solution built upon the proprietary Proteonano™ Ultraplex Proteomics Platform (autoSP3) developed by Nanomics Biotech. The autoSP3 technology comprises two core components: (1) the Proteonano™ SP3 Proteome Extract Kit, which utilizes multivalent, multi-affinity superparamagnetic nanoprobes to enable efficient protein extraction from biological samples; and (2) the Nanomation™ G1 Basic automated workstation, which delivers standardized, 96-well plate–based processing. Leveraging the autoSP3 workflow (**Fig**.1), we systematically evaluated the platform’s performance across diverse biological sample types, including cell lysates, homogenates, and trace plant tissue. Multi-center mass spectrometry analyses further demonstrated autoSP3’s robustness and translational potential, offering a reliable technological foundation to advance automated proteomic research.

**Figure 1.**
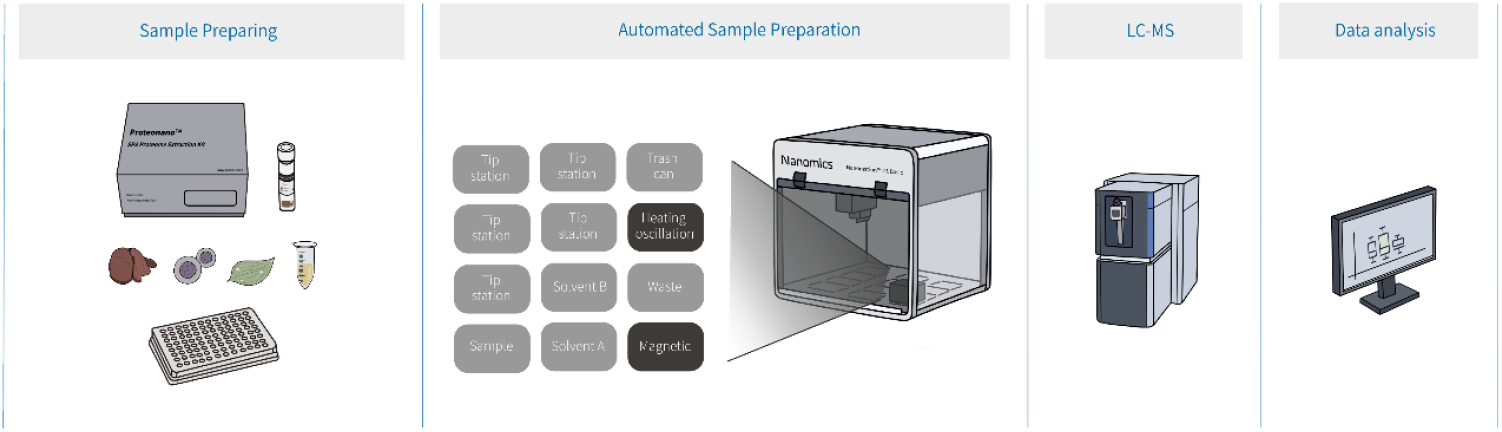
A schematic overview of the autoSP3 workflow. This diagram illustrates the complete bottom-up proteomics workflow based on autoSP3, from sample preparation through data acquisition to data analysis.

## Results

### Sample Concentration

To determine the performance of Proteonano™ SP3 Proteome Extract Kit, an analysis was performed to compare LC-MS identified protein groups from samples prepared by using the Proteonano™ SP3 Proteome Extract Kit and the organic reagent precipitation method. 10、20、50、100 and 150 μg Hela cell proteins were used for the study. LC-MS analysis was performed by using a Vanquish NEO liquid chromatography platform coupled with an Orbitrap Astral mass spectrometer with an 8 min gradient, and data was collected in DIA mode (Table 1)

**Table 1.** MS experimental parameters for different initial protein concentrations.

| MS platform | Kit | Sample | Initial sample protein amount (μg) | Injection amount (ng) | Gradient (min) |
| --- | --- | --- | --- | --- | --- |
| Orbitrap Astral | Proteonano™ SP3 Proteome Extract Kit | Hela Cell | 10 | 200 | 8 |
|  |  |  | 20 |  |  |
|  |  |  | 50 |  |  |
|  |  |  | 100 |  |  |
|  |  |  | 150 |  |  |

The Proteonano™ SP3 Proteome Extract Kit demonstrated high accuracy and data consistency when applied to HeLa cell protein samples. Peptide concentrations detected by mass spectrometry showed a linear relationship with the amount of starting protein, indicating that peptide quantification was robust across a wide range of input levels. Furthermore, the correlation coefficients of protein identification results were consistently above 0.95, confirming that the Proteonano™ SP3 Proteome Extract Kit, combined with mass spectrometry analysis, provides accurate and reproducible data under varying sample conditions (**Fig. 2**).

**Figure 2.**
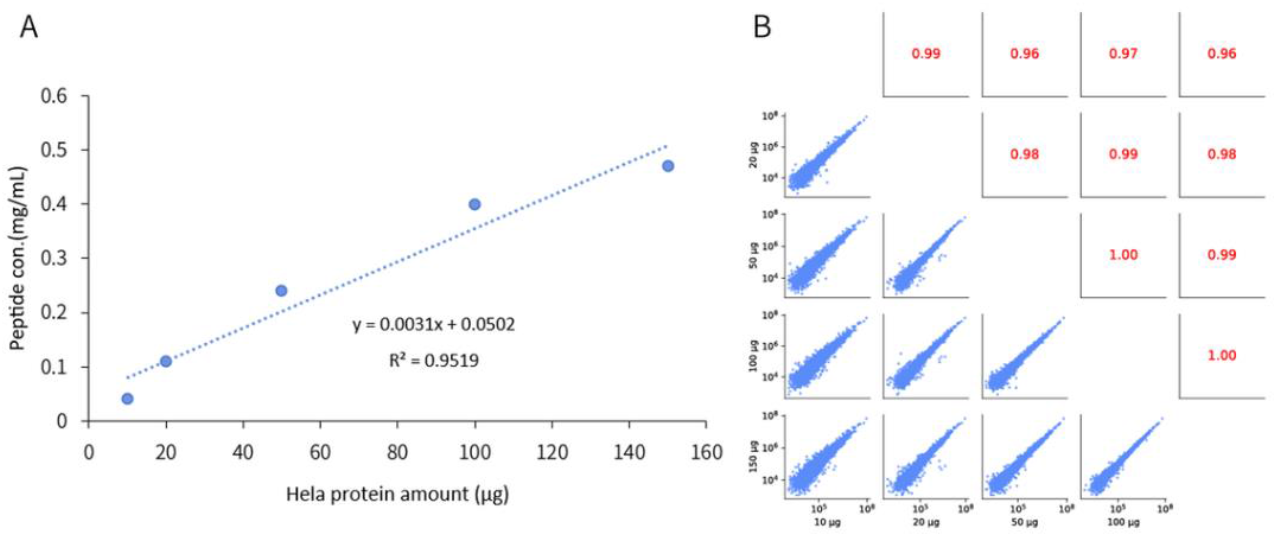
Performance of the Proteonano™ SP3 Proteome Extract Kit across varying initial protein inputs using manual sample preparation. (A) Linear range of peptide concentration in HeLa cell samples. (B) Quantitative relationship of protein concentrations in Hela cell samples.

When using the same mass spectrometry injection volume, the autoSP3 workflow showed comparable protein identification numbers to those obtained using organic reagent precipitation for HeLa samples, indicating that the Proteonano™ SP3 Proteome Extract Kit can achieve high protein identification sensitivity with minimal sample input (**Fig**. 3A). Correlation coefficients between the protein identification results of the Proteonano™ SP3 Proteome Extract Kit and the precipitation method were all above 0.90, and the average correlation coefficient between samples was 0.96, further indicating that the autoSP3 workflow has excellent accuracy (**Fig.** 3B).

**Figure 3.**
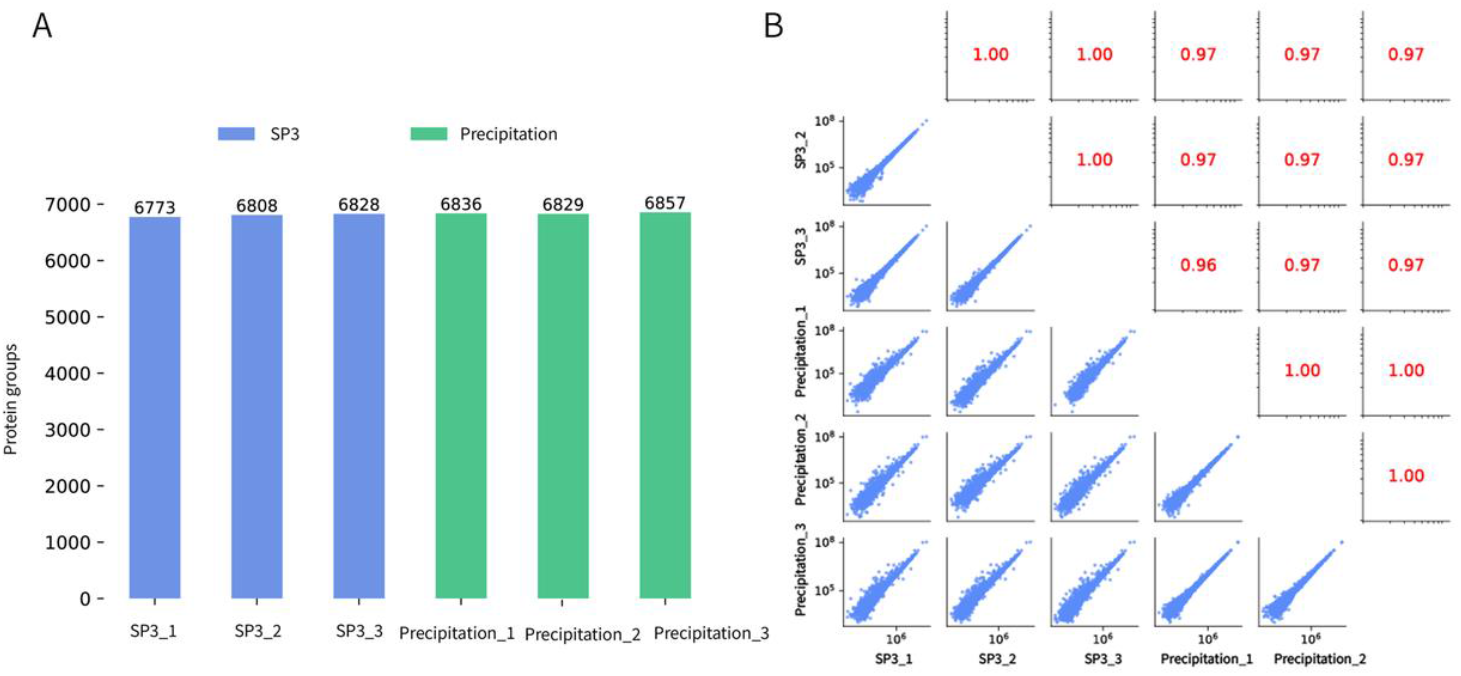
Comparison of protein data between SP3 kit and precipitation method. (A) Proteomic analysis of HeLa cell samples processed using SP3 kit (blue) and precipitation method (green). (B) Protein quantitative correlation between Proteonano™ SP3 and precipitation method.

### Comparison of Manual and Automated SP3 for Sample Preparation

Clinical proteomics faces significant challenges due to sample heterogeneity—such as fresh-frozen tissues, FFPE specimens, and blood—as well as the need for high-throughput and standardized processing. Conventional sample preparation methods are often labor-intensive, struggle with efficient detergent removal, and suffer from low recovery rates when working with limited input material. To address these limitations, the research team led by Jeroen Krijgsveld developed an automated workflow based on the SP3 method, which was applied to a cohort of 51 clinical FFPE lung adenocarcinoma (ADC) samples for proteomic analysis^3^.

Experimentally, nine mouse liver samples were processed using both manual SP3 and the autoSP3 on the Proteonano™ platform, followed by mass spectrometry. Both methods identified over 6,700 proteins, with CVs of 6.8% for manual and 6.5% for automated processing, demonstrating excellent reproducibility (**Fig**. 4A, B). Protein overlap analysis showed over 98% agreement, confirming that the Proteonano™ SP3 Kit performs equally well on the automated platform as in manual workflows (**Fig**. 4C). The autoSP3 on the Proteonano™ platform enables high-throughput, consistent, and automated sample processing, providing a reliable solution for large-scale proteomics.

**Figure 4.**
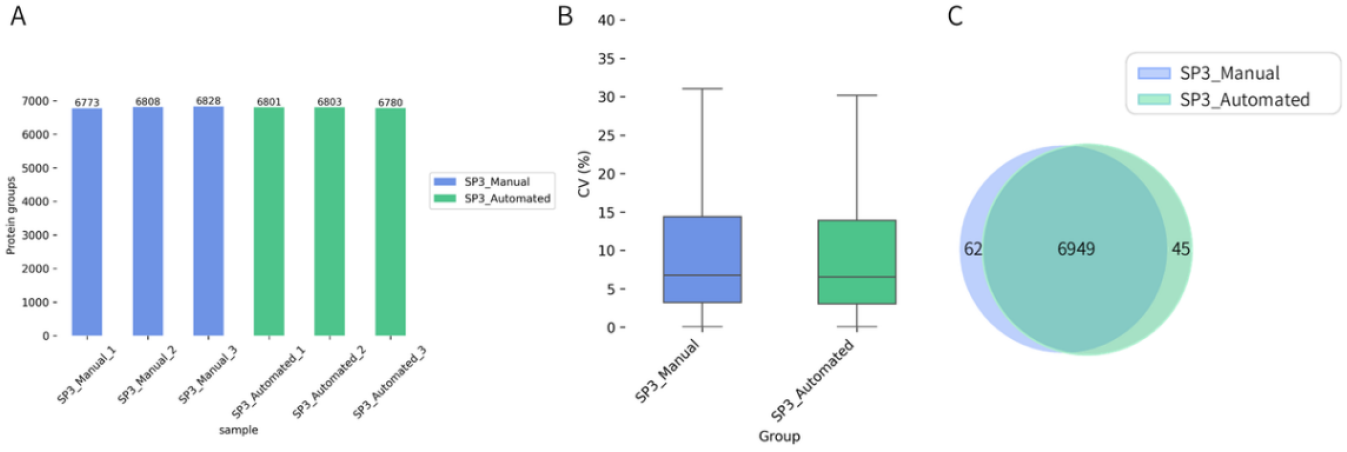
Comparison of protein data between manual and automated sample processing. (A) Proteomic analysis of mouse liver samples processed manually and with an automated platform using the SP3 kit; (B) Distribution of protein groups detected in samples processed manually and with an automated platform. (C) Overlap of protein groups identified from manual (blue) and automated (green) SP3-based sample preparation workflows. Note: Blue represents manual processing, and green represents processing with the automated platform.

### Performance of the autoSP3 on Various Biological Sample Types

Currently, various automated sample preparation methods are available to support a range of mass spectrometry (MS) acquisition strategies, including discovery-based approaches such as data-dependent acquisition (DDA-MS) ^4^ and data-independent acquisition (DIA-MS) ^5^, as well as targeted methods like parallel reaction monitoring (PRM-MS)^6^. However, automation tools and workflows must remain adaptable to meet the demands of emerging MS technologies. Developing analytical workflows that integrate automated sample preparation with LC-MS/MS—leveraging high multiplexing capability, analytical specificity, and sensitivity—is essential for advancing proteomics from basic research into clinical laboratories and precision medicine^7^.

Based on the Proteonano ™ Ultraplex Proteomics Platform, the autoSP3 can achieve deep coverage of proteomics with nanogram level protein loading. Mass spectrometry data was collected using an Orbitrap Astral high-resolution mass spectrometer. Following automatic SP3 based protein capture and tryptic protein digestion, 300 ng of peptides were aspirated by the automatic sampler and combined with the analytical column (5 μm*150 mm, C18, 2 μm, 100 Å) for separation. An 8-minute gradient was established using two mobile phases (mobile phase A: 0.1% formic acid, mobile phase B:0.1% formic acid,80% ACN). The flow rate of the liquid phase was set to 1.8 μL/min, and results were collected using Orbitrap Astral in DIA mode (Table 2). For quality control experiment, 10 μg HeLa cell protein lysate was processed by and the autoSP3 analyzed by Orbitrap Astral coupled with Vanquish NEO liquid chromatography. An 8 min gradient was used, and mass spectrometry results were collected in DIA mode.

**Table 2.**
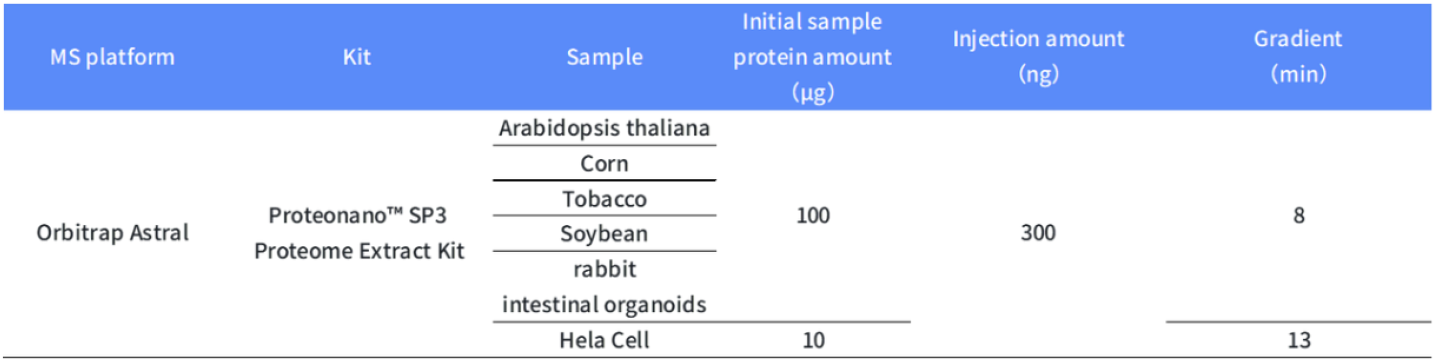
Experimental details of different biological samples.

The results demonstrate that the autoSP3 on the Proteonano™ Ultraplex Proteomics Platform enabled the identification of up to 9,414 proteins from frozen *Arabidopsis thaliana* (At.) samples. In comparison, a study by Kamil Mikulášek et al.^8^, using SP3-HILIC for protein extraction from *Arabidopsis* identified 2,979 proteins. For frozen *Tobacco* samples, the autoSP3 identified up to 9,916 proteins. Compared to commercial magnetic bead-based kits (e.g. Sera-Mag™ SpeedBeads by Cytiva), the Proteonano™ SP3 Proteome Extract Kit enabled the identification of 100 - 400 additional proteins.

Moreover, the autoSP3 identified up to 11,934 proteins from fresh *corn* samples, and up to 12,521 proteins from fresh *soybean* samples. Compared to the FASP method, the Proteonano™ SP3 Proteome Extract Kit enabled the identification of 800–1,000 additional soybean proteins (**Fig**. 5A). In a separate study, Christopher S. Hughes et al. conducted an in-depth proteomic comparison of SP3 and FASP using *yeast* whole-cell lysates (∼10 μg protein prepared in 1% SDS-containing buffer)^9^. The results showed that SP3 and FASP identified 3,944 and 4,008 proteins, respectively.

**Figure 5.**
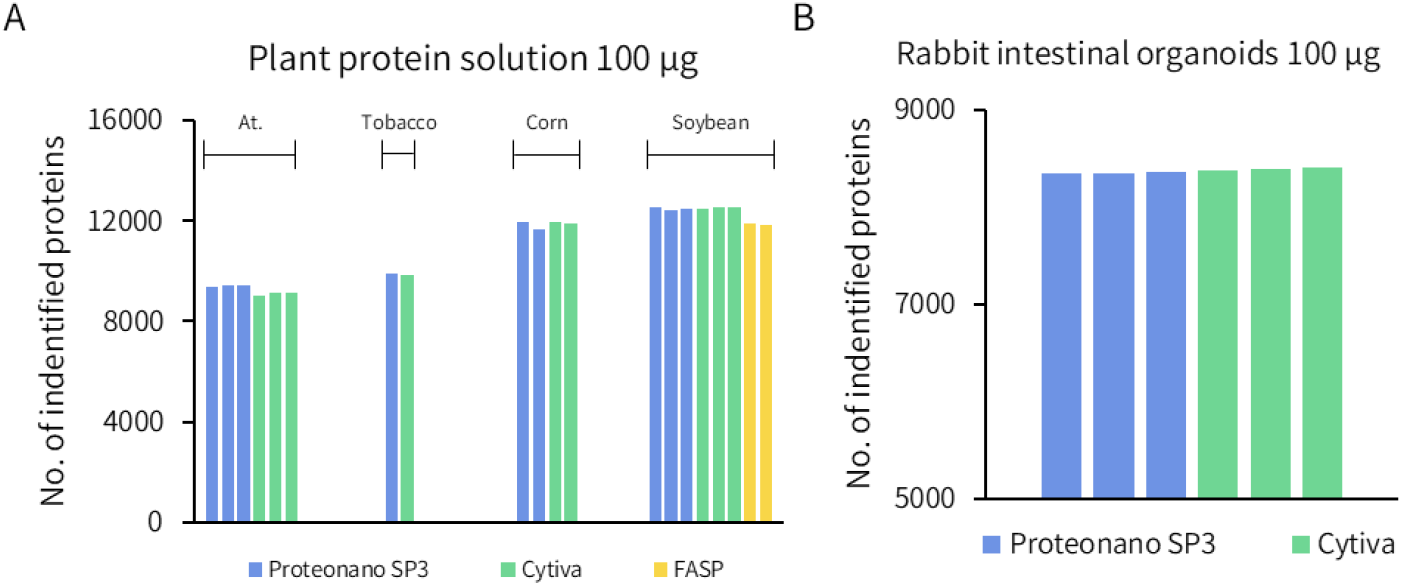
Protein identification counts for biological samples. (A) Protein identification numbers detected in plant samples processed using the Proteonano™ SP3 Proteome Extract Kit(blue), Cytiva Sera-Mag™ SpeedBeads (green), and FASP (yellow) methods. Plant samples include At, tobacco, corn, and soybean, with FASP applied only to the soybean sample. (B) Number of protein identifications detected in rabbit intestinal organoids samples processed using the Proteonano™ SP3 Proteome Extract Kit(blue) and Cytiva Sera-Mag™ SpeedBeads (green).

For frozen rabbit intestinal organoid samples, treatment with Proteonano™ SP3 Proteome Extract Kit and Proteonano™ Ultraplex Proteomics Platform combined with SP3 technology allowed detection of more than 8300 proteins, a result similar to those obtained by using the Cytiva Sera-Mag™ SpeedBeads method. Such results are highly reproducible in parallelly processed samples (**Fig**. 5B).

## Conclusion

The autoSP3 workflow based on the Proteonano™ Ultraplex Proteomics platform is designed to address sample preparation challenges in mass spectrometry. The platform combines the Proteonano™ SP3 Proteome Extract Kit and Nanomation™ G1 Basic automated workstation for high-throughput, automated protein preparation from various biological samples, including cell lysates, plant tissues, and animal tissues.

In this study, we systematically benchmarked the autoSP3 technology against traditional precipitation and FASP methods and demonstrated its superior linearity, reproducibility, and sensitivity across a broad range of input amounts. The implementation of autoSP3, integrated into the Proteonano™ Ultraplex Proteomics Platform, achieved performance equivalent to manual workflows while enabling high-throughput and standardized processing. This robust automation reduced hands-on time, minimized variability, and maintained high data quality across diverse biological matrices including cells, tissues, and plants. These results establish the automated SP3 workflow with the Proteonano Ultraplex Proteomics platform as a reliable and scalable solution for routine proteomic applications, offering strong potential to accelerate precision medicine research and clinical proteomics.

## Methods

### Reagents

Proteonano™ SP3 Proteome Extract Kit (Cat# SP3K001, Nanomics Biotech, China).

### Cell and Tissue Sources

HeLa cells were obtained from Biorun (China).

Mouse liver tissues were provided by Hangzhou Normal University (China).

### Instruments

Nanomation™ G1 Basic Automated Workstation (Nanomics Biotech, China)

Vanquish Neo UHPLC Platform (Thermo Fisher Scientific, USA)

Orbitrap Astral High-Resolution Mass Spectrometer (Thermo Fisher Scientific, USA)

### Sample Preparation

Following previous literature protocols^8^, briefly, 1 g of frozen plant tissue powder was dissolved in 1 mL of hot SDT buffer (4% SDS, 100 mM DTT, 100 mM Tris-HCl, pH 7.6). The extract was then centrifuged at 20,000 × g for 10 min at room temperature.

Animal tissue samples (∼10 mg) were placed into grinding tubes with 300 μL of 1% SDS lysis buffer containing protease inhibitors and 300 μL of ultrapure water. After vortexing, samples were incubated on ice for 1 hour, then homogenized twice at -20°C, 60 Hz for 3 minutes each. Following homogenization, nucleic acids were disrupted by sonication. Lysates were clarified by centrifugation at 15000 g for 10 minutes at 4°C, and the supernatants were collected. Protein concentrations were measured using the BCA assay.

HeLa cells were lysed directly in cell lysis buffer, and protein concentrations were determined using the BCA assay.

### Protein Extraction

For protein extraction, 100 μg of each lysate was transferred into a clean 2 mL microcentrifuge tube. Protein binding, cleanup, and enzymatic digestion were performed using the Proteonano™ SP3 Proteome Extract Kit according to the manufacturer’s protocol.

### Filter-aided sample preparation (FASP)

The FASP-based sample preparation was performed following previously published protocols^10^, primarily involving the processing of plant tissues, including reduction, alkylation, enzymatic digestion, and subsequent desalting.

### LC-MS/MS Analysis

Mass spectrometry data were collected using a liquid mass spectrometry platform with an Orbitrap Astral high resolution mass spectrometer in tandem with a Vanquish Neo liquid phase. Peptide samples were solubilized in sample buffer, aspirated by an autosampler and combined with an μPAC High Throughput column (75 μm × 5.5 cm, Thermo, USA) for separation. An 8 min analytical gradient was established using two mobile phases (mobile phase A: 0.1% formic acid and mobile phase B: 0.1% formic acid, 80% ACN). Data was acquired in data independent acquisition mode.

### Proteomic Data Analysis

The .raw files were obtained directly from mass spectrometers, or mzML files converted from .raw files by MSConvert (Version 3.0) software were searched using DIA-NN (Version 1.8.1) in library free mode. For each sample, spectra were searched against a UniProt Mouse reviewed proteome dataset, or other datasets. The DIA-NN search parameters included a 10-ppm mass tolerance for mass accuracy, one missed cleavage for trypsin, carbamidomethylation of cysteine as a fixed modification, and methionine oxidation as the sole variable modification. All other parameters were set to default, with false discovery rate (FDR) cutoffs of 0.01 applied at both the precursor and protein levels.

### Statistical Analysis and Data Visualization

The coefficient variation (CV) for each protein was determined by dividing its empirical standard deviation by its empirical mean, and CV analyses were performed on raw intensities and quantile-normalized intensities, respectively. Median values were reported as overall coefficient of variation. Pearson correlation analyses and linear regression were conducted using Pingouin^11^. No missing value imputation was performed during the above analysis, and calculations were performed only for proteins that were identified in all samples. For replicate experiments, protein identifications are expressed as AVG±SE. Seaborn was used to generate bar charts and violin charts^12^. The matplotlib-venn^13^ package and the Venn^14^ package were used to plot VENN plots.

## Acknowledgements

This work was financially supported by Nanomics Biotechnology., Ltd.

## Author contributions

H.W. and Y.W. designed the study. L.B.W. and Y.H.Z. drafted the manuscript. Y.W. led product development. L.B.W and Y.H.Z. performed the proteomics data analysis. X.H.O. conducted the sample preparation and mass spectrometry experiments.

